# Human influences on antipredator behaviour in Darwin’s finches

**DOI:** 10.1101/591651

**Authors:** Kiyoko M. Gotanda

## Abstract

1) Humans exert dramatic influences upon the environment, creating novel selective pressures to which organisms must adapt. On the Galapagos, humans have established a permanent presence and have altered selective pressures through influences such as invasive predators and urbanization, affecting iconic species such as Darwin’s finches.

2) Here, I ask two key questions: (i) does antipredator behaviour (e.g. FID) change depending on whether invasive predators are historically absent, present, or eradicated? and (ii) to what degree does urbanization affect antipredator behaviour? This study is one of the first to quantify antipredator behaviour in endemic species *after* the eradication of invasive predators. This will help to understand the consequences of invasive predator eradication and inform conservation measures.

3) I quantified flight initiation distance (FID), an antipredator behaviour, in Darwin’s finches, across multiple islands in the Galapagos that varied in the presence, absence, or successful eradication of invasive predators. On islands with human populations, I quantified FID in urban and non-urban populations of finches.

4) FID was higher on islands with invasive predators compared to islands with no predators. On islands from which invasive predators were eradicated ∼11 years previously, FID was also higher than on islands with no invasive predators. Within islands that had both urban and non-urban populations of finches, FID was lower in urban finch populations, but only above a threshold human population size. FID in larger urban areas on islands with invasive predators was similar to or *lower* than FID on islands with no history of invasive predators.

5) Overall, these results suggest that invasive predators can have a lasting effect on antipredator behaviour, even after eradication. Furthermore, the effect of urbanization can strongly oppose the effect of invasive predators, reducing antipredator behaviour to levels lower than found on pristine islands with no human influences. These results improve our understanding of human influences on antipredator behaviour which can help inform future conservation and management efforts on islands.

## Introduction

> “All of [the terrestrial birds] are often approached sufficiently near to be killed with a switch, and sometimes, as I myself tried, with a cap or a hat.” – Charles Darwin in “The Voyage of the Beagle”

Human influences such as invasive species and urbanization can strongly affect the process of local adaptation (Alberti et al., 2017; Johnson & Munshi-South, 2017; Low, 2002; Vitousek, Mooney, Lubchenco, & Melillo, 1997). Such effects are amplified on islands such as the Galapagos Islands, where small population sizes and strong isolation increase the vulnerability of local flora and fauna to human influences, often resulting in loss of island biodiversity through extinctions (Blackburn, Cassey, Duncan, Evans, & Gaston, 2004; Fritts & Rodda, 1998; MacArthur & Wilson, 1967; Sax & Gaines, 2008). Among the endemic species on the Galapagos Islands are Darwin’s finches, an iconic example of an adaptive radiation in which a single founding species has evolved into several species, each with different adaptions (e.g. beak shapes and body sizes) to exploit different ecological niches (Grant & Grant, 2008, 2014). Humans began establishing settlements on the Galapagos in the early 19^th^ century (Fitter, Fitter, & Hosking, 2016), and since then, human influences such as invasive predators and urbanization have affected several islands on the Galapagos. Many organisms initially respond to such human influences through behavioural adaptations. Here, I consider how two human influences – invasive predators and urbanization – might alter antipredator behaviour in Darwin’s finches on the Galapagos Islands.

Invasive predators have strong ecological and evolutionary effects (Clavero, Brotons, Pons, & Sol, 2009; Lever, 1994; Low, 2002; Vitousek et al., 1997), and this impact is known to be correlated with local extinction events (Clavero & García-Berthou, 2005; McKinney & Lockwood, 1999). On islands, the lack of predators and correlated relaxed selection can result in reduced antipredator behaviour (Beauchamp, 2004; Blumstein, Daniel, & Springett, 2004; Blumstein, 2002; Jolly, Webb, & Phillips, 2018). This evolutionary naïveté of isolated animals that have evolved without major predators can contribute to the extirpation of island species (Blumstein et al., 2004; Blumstein, 2002; Blumstein & Daniel, 2005; Cooper, Pyron, & Garland, 2014). In particular, feral and domestic house cats (*Felis silvestris catus*) are of concern for island biodiversity because cats target small animals such as birds and reptiles (Blancher, 2013; Loss, Will, & Marra, 2013; Woinarski et al., 2018; Woods, Mcdonald, & Har Ris, 2003), and invasive house cats now exist on four islands of the Galapagos (Phillips, Wiedenfeld, & Snell, 2012), presenting a critical threat for Galapagos biodiversity (Stone, Snell, & Snell, 1994; Wiedenfeld & Jiménez-Uzcátegui, 2008). Past research on the effects of invasive predators in the Galapagos has focused on behavioural adaptations in reptiles (e.g. Berger, Wikelski, Romero, Kalko, & Roedl, 2007; Stone et al., 1994), and thus, little is known about the effect of novel mammalian predators on endemic land birds. Given the resulting selective pressures, natural selection should favour an increase in antipredator behaviour after the introduction of an invasive predator to reduce mortality.

Effective conservation management, especially on islands, often involve eradication of invasive predators to protect the local and endemic species (Jones et al., 2016; Nogales et al., 2013). Post-eradication research typically follows local and endemic species population recovery (Côté & Sutherland, 1997; Hughes, Martin, & Reynolds, 2008; Lavers, Wilcox, & Donlan, 2010), monitors the re-introduction of extirpated species to previously abandoned breeding grounds (Campbell et al., 2011), or focus on major ecological effects such as changes in food web dynamics (Hughes et al., 2008; Zavaleta, Hobbs, & Mooney, 2001). All this research contributes to the growing need to understand post-eradication effects (Côté & Sutherland, 1997; Lavers et al., 2010; Phillips, 2010), yet surprisingly little research has focused on post-eradication behavioural adaptations, nor how quickly such behavioural adaptations might occur. Post-eradication behavioural adaptations could have population-level consequences on fitness. For example, increased antipredator behaviour can have associated costs due to the reallocation of energy and time away from other important behaviours such as foraging, reproduction, and rearing of young (Cooper & Frederick, 2007; Cooper & Blumstein, 2015; Ydenberg & Dill, 1986). Thus, if increased antipredator behaviour is maintained after eradication, this might result in a decrease in fitness for local and endemic species. Understanding how local and endemic species will behaviourally adapt post-eradication could help improve conservation efforts. On the Galapagos, some islands have invasive house cats, some have remained free of invasive predators, and some islands have successfully eradicated invasive predators. This allows for among-island comparisons of antipredator behaviour in relation to the current and historical invasive-predator regime.

Urbanization has rapidly increased in the past century, with more than half the world’s population occupying urban settlements, severely altering patterns of selection and adaptation (Alberti et al., 2017; Hendry, Gotanda, & Svensson, 2017; Johnson & Munshi-South, 2017). In general, animals such as birds show decreased antipredator behaviour in urban areas compared to rural areas, likely due to habituation to humans (Díaz et al., 2013; Møller, 2009; Møller & Tryjanowski, 2014; Samia et al., 2017). However, such a reduction in antipredator behaviour in urban areas could make organisms more vulnerable to different threats, such as invasive predators. Quantifying the degree to which urbanization can reduce antipredator behaviours can inform our understanding of the impacts of urbanization. Can urbanization reduce antipredator behaviour to levels *before* the introduction of predators?

The Galapagos Islands represent an excellent opportunity to study the effects on invasive predators and urbanization for two key reasons that few, if any, other systems offer. First, few archipelagos in the world have islands that vary not only in the presence or absence of invasive predators, but also have islands that have successfully eradicated invasive predators. Second, islands differ not only in the presence or absence of urban centres, but also in the size of the urban centres, representing a novel opportunity to compare antipredator behaviour along a gradient of urbanization as well as among islands that differ in the presence or absence of urban centres. The isolation of the Galapagos Islands removes potentially confounding factors such as high gene flow or continued influxes of introduced predators, allowing me to ask two key questions regarding human influences and antipredator behaviour. First, I ask how will antipredator behaviour change depending on whether invasive predators are present, absent, or perhaps more importantly, eradicated from an island? Very little research has been done on behavioural adaptations in endemic species post invasive predator eradication. Second, I ask how much can urbanization reduce antipredator behaviour – can it be reduced to levels found on islands with no history of invasive predators? Together, these two questions can inform how human influences are affecting antipredator behaviour on isolated islands.

## Materials and methods

### Site descriptions

The Galapagos Islands are a volcanic archipelago located ∼1,000 km off the coast of Ecuador. Local or endemic predators such as owls or snakes are found on all islands (Supplemental Table 1; Swash & Still 2005). Snakes are thought to prey on the nestlings of ground finches and are thus an unlikely predator. However, short-eared owls, found on all islands surveyed in this study, are known predators of adult land birds such as Darwin’s finches (Groot, 1982). Galapagos Hawks, also predators of ground finches (Vries, 1976, 2015), are found on four (Santa Fe, Española, Isabela, and Santa Cruz) of the eight islands surveyed in this study (Swash & Still, 2005; Vries, 1989). Unfortunately, little data are available about the current densities of local and endemic predators on these islands; however, the ecology of the local and endemic predators (e.g. owls) is well documented and can thus be assumed to be predators of finches. The invasive-predator regime (presence of house cats) on the islands (Supplemental Table 1) were classified as: present (Floreana, Isabela, San Cristobal, Santa Cruz), pristine (Santa Fe, Española), or successful eradication (Baltra, North Seymour). House cats and rodents were successfully eradicated from Baltra in 2003 (R. B. Phillips et al., 2005; R.Brand Phillips et al., 2012), and rats from North Seymour in 2008 (Carrión, Sevilla, & Tapia, 2008; Harper & Carrión, 2011); I will refer to these as “eradicated” below for brevity. The two pristine islands and two eradicated islands have no permanent human populations. On the four islands with human populations (Floreana, Isabela, San Cristobal, Santa Cruz), site urbanization categories were classified as: urban (in town) or non-urban (remote areas several kilometres away from town and not visited by tourists). No islands with permanent human populations and no presence of invasive predators nor pristine islands with permanent human populations exist in the Galapagos archipelago, and thus, I am restricted to the among island comparisons outlined above.

### Data collection

Flight initiation distance (FID), the distance at which a prey flees an approaching predator, is a metric used to quantify antipredator behaviour (Cooper & Frederick, 2007; Cooper & Blumstein, 2015; Ydenberg & Dill, 1986). An individual’s decision to flee is influenced by the perceived costs and benefits of remaining or taking flight, which means FID can be an indicator of how an organism assesses risk, and thus, antipredator behaviour. Data were collected from 2015 to 2018, generally between February and April (some data on San Cristobal were collected in November 2017), on eight islands of the Galapagos archipelago: four islands with the presence of invasive predators, two islands with no history of invasive predators, and two island that have successfully eradicated invasive predators (Supplemental Figure 1). Due to finch species distributions varying across islands as well as Darwin’s finch’s species being closely related, data from the only species that occurs on all islands, the small ground finch, *Geospiza fuliginosa*, are presented.

FID measurements were performed with a human stimulus following methods from Blumstein (2006). A focal finch was located by walking and searching the landscape at a slow walking pace, and the finch’s initial behaviour was noted. To minimize the possibility of pseudoreplication, in a given day, each trial ensured the focal finch was of a different sex or age class (for males) than finches that had previously been approached. However, it is possible the same bird might have been approached on different days or years because the finches were not individually banded. Birds were located in areas that had relatively open habitat to ensure a straight approach by the human and a clear sightline from the human to the finch. The human would then approach the focal finch at a standardized speed (∼0.85 m/s). Human stimuli always wore neutral-coloured clothing and looked at the focal individual while approaching.

Flight was considered to have been initiated if the finch extended its wings and flew; the distance flown could be short (<0.5 m) or substantial (out of sight). Finches that hopped away instead of taking flight were omitted from the study, though this was an extremely rare occurrence (two or three occurrences across all sampling periods). A marker was placed where the finch originally was and where the stimulus was when the finch took flight, and the distance between these markers was the FID. Because of the complexity of the landscape, the distance at which the stimulus started from could not be standardized (Blumstein, 2003, 2006; Samia, Nomura, & Blumstein, 2013), and so I noted the distance from where the human started to the flight-initiation marker (starting distance). Alert distance, the distance at which an individual is aware of the approaching stimulus (Cooper, Samia, & Blumstein, 2015), could not be quantified because the focal individual was often foraging on the ground and would repeatedly raise its head, and so normal foraging behaviour was indistinguishable from an alert reaction to an approaching human.

Each data point collected included the island, invasive-predator regime, urbanization category (on islands with permanent human populations), sex, time of day, and group size (defined as at least one other finch within a 1m radius of the focal finch and if so, how many other finches were within 1m of each other). A minimum of six finches were sampled at each site (Supplementary Table 1). Finch sex was identified by plumage, and for ambiguous cases sex was denoted as unknown. Time of day was noted because birds are most active at dawn and at dusk, so baseline activity levels and behaviours can vary throughout the day. Group size was noted because it could increase FID because larger groups mean more observers and thus, detection of a potential threat will occur when the threat is still a longer distance away (Ydenberg & Dill, 1986). Conversely, group size could decrease FID through the dilution effect where the probability a predator will target a specific individual decreases as group size increases (Cooper & Blumstein, 2015).

### Statistical analysis

All analyses were done in R (version 3.4.3). To meet assumptions of normal distributions, FID and starting distance (the distance between the focal finch and the stimulus starting position) were log-transformed. Then, FID and starting distance were centered by subtracting the mean from each value and then dividing by the standard deviation. Because starting distance could not be standardized and is known to affect FID, it was included in the analyses, and group size and time of day were included as covariates. Lastly, sex was included as a fixed factor and island was included as a random factor as appropriate. Further analysis details are below, but in general, linear mixed models were performed using lmer() from the lme4 package (Bates, Maechler, Bolker, & Walker, 2015) and Anova() from the car package (Fox & Weisberg, 2011). Random-factor significance was determined with ranova(), and post-hoc pairwise comparisons for fixed factors in the linear mixed models used difflsmeans(), both from the lmerTest package (Kuznetsova, Brockhoff, & Christensen, 2017). R^2^ values for mixed models were calculated with the r2beta() from the r2glmm package (version 0.1.2) and η^2^ values for linear models were calculated with etaSquared() from the lsr package (version 0.5).

### Does antipredator vary depending on whether invasive predators are historically absent, present, or have been eradicated?

On islands that had both urban and non-urban populations, only data from non-urban sites were used for this analysis. For this analysis, the number of finches sampled on each island ranged from 9 finches to 29 finches. A linear mixed model was first performed with invasive-predator regime, sex, and starting distance (and their interactions) as fixed factors, island a as random factor, and time of day and group size as covariates. None of the two- or three-way interactions were significant (0.634 < P < 0.924) and the best-fit model had the interactions removed. Post-hoc comparisons focused on pairwise comparisons between invasive-predator regimes.

To determine whether FID varied among islands within a given invasive-predator regime, I ran a linear model with island, sex, and starting distance (and their interactions) as fixed factors and time of day and group size as covariates. The only significant interaction was between island and starting distance. The best-fit model had the non-significant interactions removed. Post-hoc comparisons were focused on pairwise comparisons between islands within a given invasive-predator regime. A Tukey’s HSD (honestly significant difference) test was used for post-hoc comparisons for this analysis.

### How much can urbanization affect antipredator behaviour?

Only data collected from the four islands with permanent populations were used for this analysis. For this analysis, the number of finches sampled in each population ranged from 6 finches to 29 finches. A linear mixed model was performed with site urbanization, sex, and starting distance (and their interactions) as fixed factors, island as a random factor, and time of day and group size as covariates. None of the two- or three-way interactions were significant (0.244 < P < 0.560) and model with no interactions was a better fit and is reported here. To look at within island differences in FID among urban and non-urban finches, for each island, a linear model was performed with the same factors as above without island as a random factor.

## Results

### Does antipredator vary depending on whether invasive predators are historically absent, present, or have been eradicated?

On islands with invasive predators and islands where predators have been eradicated, FID in finches was significantly higher than on pristine islands (Table 1, Figure 1). Post-hoc comparisons showed that finches on pristine islands (Santa Fe and Española) had lower FID when compared to finches on islands with invasive predators (Figure 1; p = 0.009) and eradicated islands (Figure 1; p = 0.005), and finches on eradicated islands did not differ in FID when compared to finches on islands with invasive predators (Figure 1, p = 0.353). When considering the effect of island independently of invasive-predator regime, island had a significant effect on FID (Figure 1; Table 2). A significant interaction between island and starting distance was found (Supplemental Figure 2; Table 2). Post-hoc comparisons of FID within an invasive-predator regime showed no significant difference in FID between finches on the two pristine islands (p = 1.000), among the four islands with invasive predators (0.458 < p < 1.000), or between the two eradicated islands (p = 0.994).

**Figure 1.**
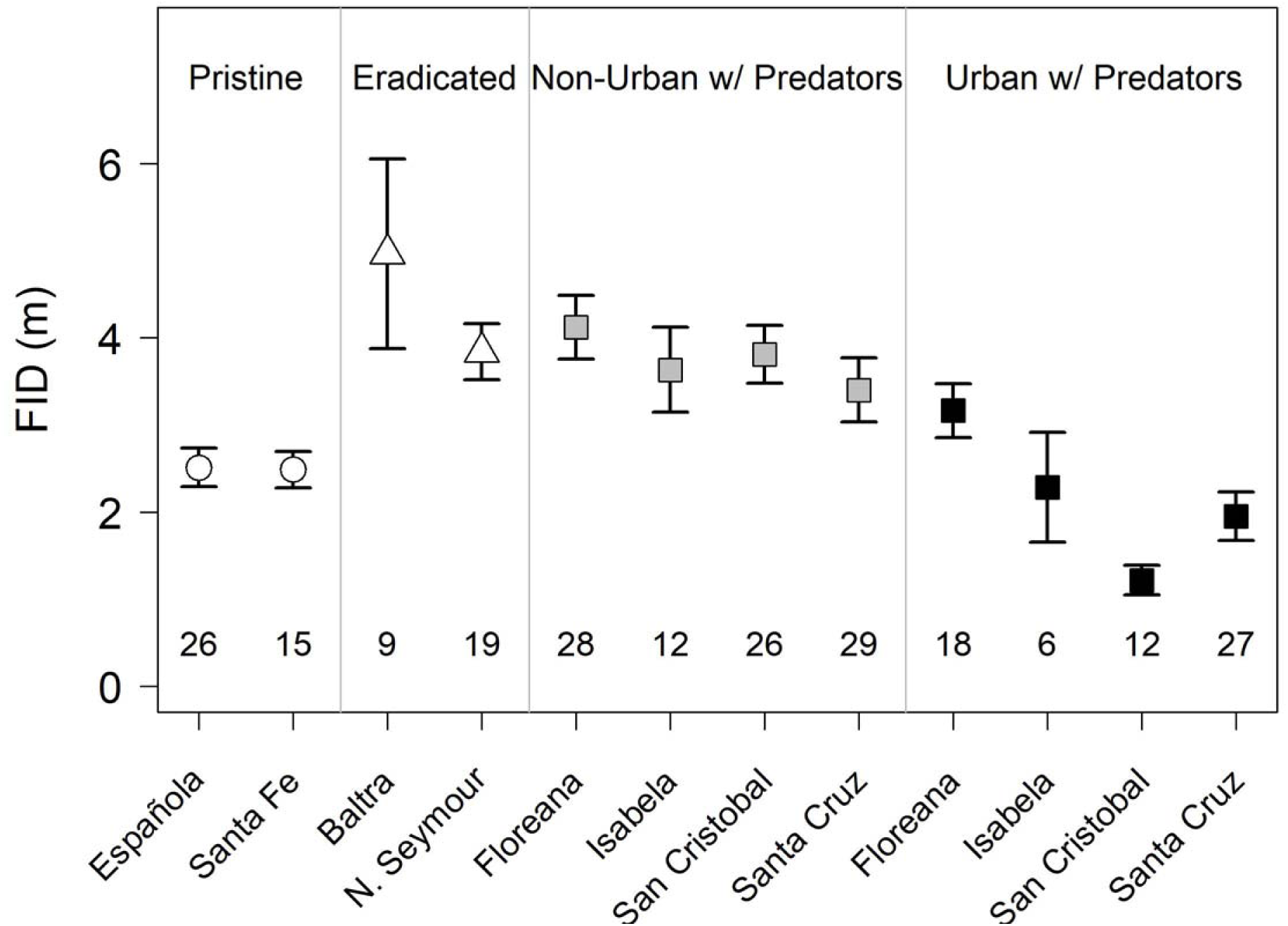
Mean FID and standard error on the eight islands data were collected from. Symbol shape denotes the invasive-predator regime (pristine, eradicated, or present). Island and site categorization are listed on the top, and numbers indicate sample sizes. Symbol colour indicates the site urbanization category on the four islands with permanent human populations and invasive predators (Floreana, Isabela, San Cristobal, and Santa Cruz). Grey colour indicates non-urban finches, and black indicates urban finches. Open symbols indicate islands with no permanent human populations. The two far left islands, Española and Santa Fe are pristine islands with no history of invasive predators. The next two islands, Baltra and North Seymour are where invasive predators have been successfully eradicated in 2003 and 2008 respectively, and show maintained increased antipredator behaviour. Islands with predators are ordered by population size with Floreana having the smallest urban population and Santa Cruz the largest urban population. In non-urban sites that have invasive predators, antipredator behaviour is increased, and in urban sites, antipredator is significantly decreased on all islands except Floreana. On San Cristobal and Santa Cruz, the antipredator behaviour is lower than on islands untouched by invasive predators and humans.

**Table 1.** Results looking at the effects of invasive predator regime (present, pristine, or eradicated) on flight initiation distance (FID). A general linear mixed model was performed with FID as the dependent variable, invasive-predator regime, sex, and starting distance as fixed factors, time of day and group size as covariates, and island as a random factor (denoted by *). Non-significant interactions were removed from the analysis. R^2^ for the full model was 0.142. Data used for this analysis were from eight islands that varied in invasive-predator regime. For data collected on islands with permanent human populations, only data from non-urban sites used here. Bold indicates significant P values.

| | $\chi^2$ | d.f. | $R^2$ | P |
| --- | --- | --- | --- | --- |
| Predation | 16.770 | 2 | 0.093 | **<0.001** |
| Sex | 3.505 | 2 | 0.008 | 0.173 |
| Time of Day | 1.215 | 1 | 0.007 | 0.270 |
| Starting Distance | 0.004 | 1 | 0.000 | 0.949 |
| Group Size | 0.207 | 1 | 0.010 | 0.207 |
| Island* | 0.182 | 1 | NA | 0.669 |

**Table 2.** Results looking at the effect of island on flight initiation distance (FID). A linear mixed model was performed with FID as the dependent variable, island, sex, and starting distance as fixed factors and time of day and group size as covariates. Non-significant interactions were removed from the analysis. Data used for this analysis were from eight islands that varied in invasive-predator regime. On islands with human populations, only data from non-urban sites are included. Bold indicates significant P values.

| | F | d.f. | $\eta^2$ | P |
| --- | --- | --- | --- | --- |
| Island | 4.087 | 7 | 0.140 | **<0.001** |
| Sex | 1.832 | 2 | 0.018 | 0.164 |
| Time of Day | 2.204 | 1 | 0.011 | 0.140 |
| Starting Distance | 0.330 | 1 | 0.000 | 0.567 |
| Group Size | 3.862 | 1 | 0.019 | 0.051 |
| Island * Starting Distance | 3.522 | 7 | 0.120 | **0.002** |

### How much can urbanization affect antipredator behaviour?

Urban finches had significantly lower FID as compared to non-urban finches (Table 3, Figs. 1 and 2). A post-hoc linear model analysing FID in finches on Floreana showed no significant difference between urban and non-urban populations (F = 0.158, p = 0.514) whereas FID in finches was significantly lower in urban areas as compared to non-urban areas on Isabela (F = 9.005, p = 0.011), San Cristobal (F = 28.000, p < 0.001), and Santa Cruz (F = 7.555, p = 0.008). On San Cristobal and Santa Cruz, finches in urban populations had lower FID than found on pristine islands (Figure 1). Group size was positively correlated with FID (Table 3, Supplemental Figure 3) for both urban and non-urban finches.

**Figure 2.**
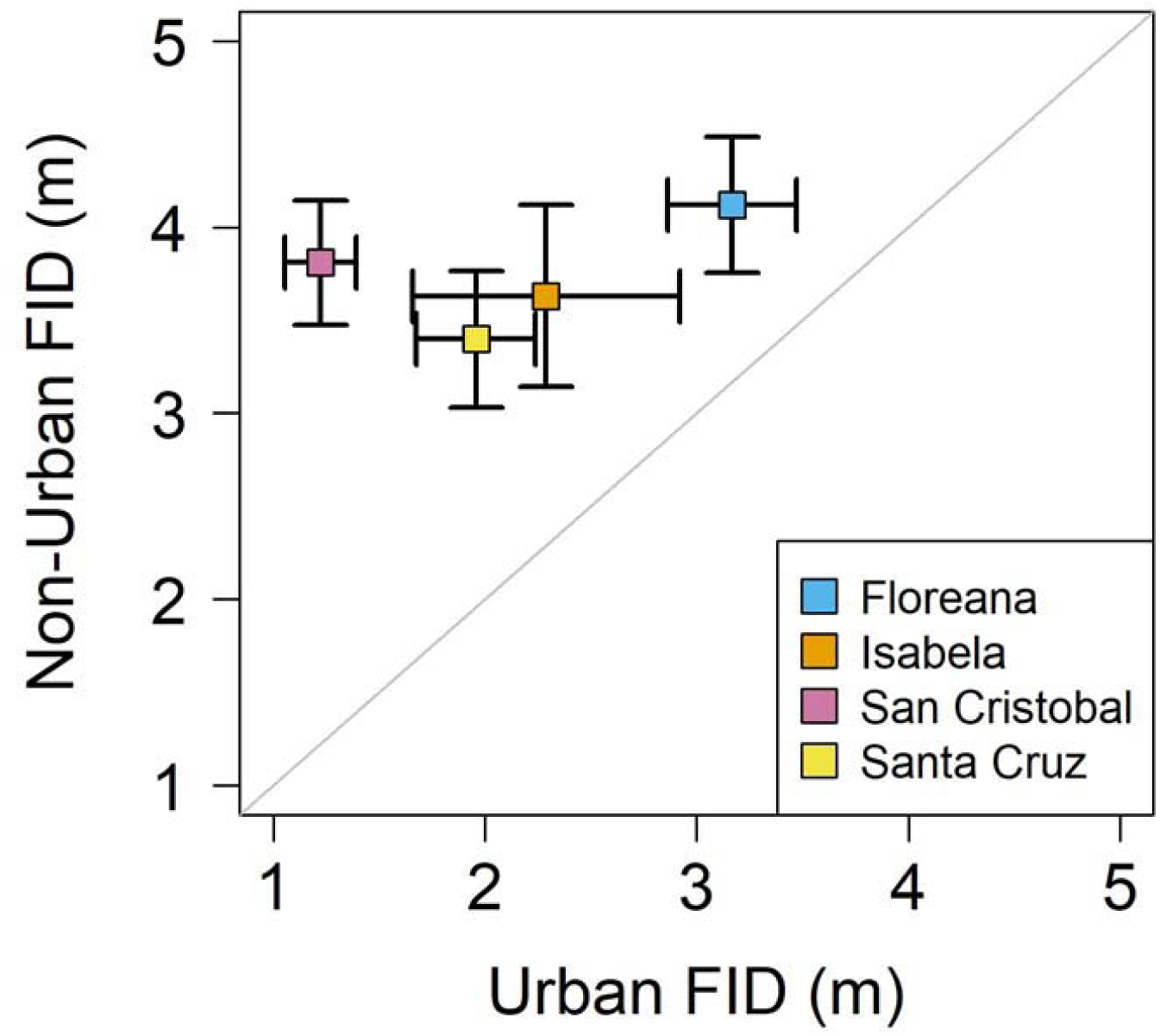
Mean FID and standard error contrasting urban finches with non-urban finches from the four islands that have permanent human populations. Colour indicates island. Above the 1:1 line indicates FID in non-urban areas is higher than FID in urban areas. Floreana, the island with the smallest human population size, has the least difference in FID between urban and non-urban finches.

**Table 3.** Results looking at the effect of urbanization on flight initiation distance (FID). A general linear mixed model was performed with FID as the dependent variable, site urbanization category, sex, and starting distance as fixed factors, time of day and group size as covariates, and island as a random factor (denoted by *). Non-significant interactions were removed from the analysis. R^2^ for full model was 0.258. Data used in this analysis were from urban and non-urban sites on the four islands with permanent human populations. Bold indicates significant P values.

| | $\chi^2$ | d.f. | $R^2$ | P |
| --- | --- | --- | --- | --- |
| Site Category | 43.307 | 1 | 0.215 | **<0.001** |
| Sex | 0.041 | 1 | 0.033 | 0.839 |
| Time of Day | 2.371 | 1 | 0.015 | 0.124 |
| Starting Distance | 2.000 | 1 | 0.069 | 0.157 |
| Group Size | 5.420 | 1 | 0.033 | **0.020** |
| Island* | 2.284 | 1 | NA | 0.131 |

## Discussion

### Increased antipredator behaviour is maintained after eradication of invasive predators

Finches exhibited increased antipredator behaviour on islands with invasive predators (Table 1; Figure 1). More interestingly, this increased antipredator behaviour was also observed on islands where invasive predators had been eradicated (in 2003 and 2008, 13 and 8 years prior to data collection; the mean generation time of finches is one year, by comparison). Since all islands have naturally occurring local or endemic predators of finches (Supplemental Table 1), this observation was most likely due to the presence of invasive predators, even after eradication. This is one of the first studies to show that increased antipredator behaviour has been maintained on islands that have had invasive predators removed. Several possible reasons exist for these observations, especially the apparent maintenance of elevated FID on eradicated islands. First, it is possible increased antipredator behaviour has evolved on islands that have and used to have invasive predators. However, without knowledge of heritability, and thus actual evolution, this cannot be confirmed, but would be an area for future research. Second, perhaps the expected costs of increased antipredator behaviour are not high enough to cause a reversion to pre-predator levels. Increased FID can have associated costs (Cooper & Frederick, 2007; Cooper & Blumstein, 2015; Ydenberg & Dill, 1986), suggesting that if this behavioural adaptation were costly then finches on eradicated islands would have FID comparable to finches on pristine islands. It could also be that not enough time has elapsed for reversion in antipredator behaviour. Third, the multi-predator hypothesis (Blumstein, 2006) suggests that increased antipredator behaviour can be maintained by the presence of remaining local and endemic predators. However, the lower antipredator behaviour on islands with no history of invasive predators suggest that it is the invasive predators that have caused the increase in antipredator behaviour, and a different mechanism is maintaining the increased antipredator behaviour. Fourth, cultural transmission of increased FID (Aplin et al., 2015; Aplin, 2018) with learned behaviour transmitted from generation to generation could maintain the increased FID. Lastly, the increase in FID could be due to something other than predation (e.g. parasitism, life history, or habitat; Cooper & Blumstein, 2015), which could still be present on eradicated islands, or that for unknown historical reasons, antipredator behaviour on the eradicated islands have historically been high. The last reason is possible, but it would be a quite a coincidence if the eradicated islands that have this elevated FID for some reason unrelated to predation just happen to be exactly the islands where predation was introduced and then eradicated.

Regardless of the mechanism, the fact that antipredator behaviour levels did not revert post-eradication (when comparing FID on eradicated islands to FID on pristine islands; Figure 1) has potential consequences for evaluating the efficacy of eradication efforts. Recent studies of local animal populations post-eradication have focused on demographic parameters such as population recovery (Côté & Sutherland, 1997; Hughes et al., 2008; Lavers et al., 2010) or on ecological parameters such as food-web dynamics (Hughes et al., 2008; Zavaleta et al., 2001). However, such phenomena will be influenced by behavioural shifts. For example, increased antipredator behaviour correlates with decreased time and energy for behaviours such as foraging, courting, defending territories, or caring for offspring (Cooper & Frederick, 2007; Cooper & Blumstein, 2015; Ydenberg & Dill, 1986), which could affect population recovery and/or food-web dynamics. Thus, understanding how the eradication of invasive predators will affect the behaviour of local or endemic animals should be central to future conservation efforts (Côté & Sutherland, 1997; Lavers et al., 2010; R. A. Phillips, 2010).

### Urbanization can decrease antipredator behaviour to levels lower than before the introduction of predators

Finches in urban areas had lower FID than finches in non-urban areas, supporting previous findings (Díaz et al., 2013; Møller, 2009; Møller & Tryjanowski, 2014; Samia et al., 2017). However, two interesting points are found in this general trend. First, the *degree* of urbanization appears to determine just how much lower FID is for urban finches when compared to non-urban finches. Finches in the town of Puerto Velasco Ibarra on Floreana had the highest FID compared to finches in other towns, and Puerto Velasco Ibarra is also the smallest town, with a permanent population of only 111 (Supplemental Table 1). The significantly lower FID of finches in larger towns as compared to non-urban finches suggests an urbanization “threshold”, such that the degree of urbanization needs to be high enough to exert sufficient selective pressure on finches to drive behavioural adaptation; in short, perhaps Puerto Velasco Ibarra is simply too small to be an “urban” site for the purposes of finch behavioural adaptation. The next largest town, Puerto Villamil on Isabela, with a population of 2,164 (Supplemental Table 1), had significantly lower FID than Puerto Velasco Ibarra on Floreana. Puerto Villamil is still a relatively small town (Supplemental Table 1), showing that the threshold amount of urbanization sufficient to produce differences in antipredator behaviour is not very high.

The second interesting point is that on some islands, urbanization can result in FID that is *lower* than FID on islands that have never been exposed to predators (Figure 1), even though urban areas invariably contain invasive predators such as cats and rats. In other words, in some towns, such as Puerto Baquizo Moreno on San Cristobal and Puerto Ayora on Santa Cruz, FID has been reduced to levels below what was observed on islands with no history of invasive predators. Such reductions in FID in urban finches is likely due initially to habituation (Blumstein, Fernandez-Juricic, Zollner, & Garity, 2005; Cavalli, Baladrón, Isacch, Biondi, & Bó, 2018; Møller, 2008; Samia, Nakagawa, Nomura, Rangel, & Blumstein, 2016) which can lead to lower FID being an evolved adaptation (van Dongen, Robinson, Weston, Mulder, & Guay, 2015) This suggests pressures from urbanization on antipredator behaviour can be so strong that it results in FID lower than the baseline FID quantified on pristine islands, counteracting any increase in FID due to the presence of invasive predators. For both this and the previous point, more research needs to be conducted on more systems with town of varying size given the relatively small number of islands and populations in this study. Altogether, this suggests that the effects of urbanization on organisms can be quite strong, with likely evolutionary and ecological consequences (Alberti, 2015; Alberti et al., 2017; Johnson & Munshi-South, 2017).

### Group size and starting distance

FID significantly increased with increasing finch group size for urban and non-urban populations on islands with permanent human populations (Supplemental Figure 3). This supports the “many-eyes hypothesis” that detection of predators occurs earlier in large groups, when the predator is further away, due to the larger number of individuals watching (Ydenberg & Dill, 1986). However, an effect of group size on FID in relation to invasive predation regime was not found, indicating that on islands with no history of or having successfully eradicated invasive predators, group size does not affect FID in individual finches. Perhaps the vigilance achieved with “many-eyes” is necessary on islands with predators, but not in the absence (historically or eradicated) of invasive predators.

Interestingly, starting distance was not significantly positively correlated with FID as expected based on previous research (Blumstein, 2003; Samia et al., 2013). In fact, on one island, San Cristobal, the relationship was negative with longer starting distances resulting in shorter FIDs (Supplementary Figure 2), and this relationship could be explored with further research. However, the effect of starting distance on FID is highly variable among species and a some species do not have a positive correlation between starting distance and FID (Blumstein, 2003). Thus. it is possible that Darwin’s finches are one of the very few species where starting distance has a very low impact on FID and that is why a positive correlation between starting distance and FID was not detected in this study.

### Conclusions

The Galapagos Islands represent an opportune system to study the effects of human influences due to the among island differences in invasive-predator regime as well as the differences in the amount of urbanization. Such systems do not readily exist elsewhere. Here, I showed how antipredator behaviour increased in response to invasive predators and was maintained even after eradication of those invasive predators, and this can have possible demographic and ecological effects. This is one of the first studies to look at post-eradication behaviours in local and endemic species, which can have implications for future conservation efforts. I also found that urbanization can reduce antipredator behaviour to levels at or below what was found on pristine islands, attesting to the strength of the effect urbanization can have on behavioural traits. Understanding the effects of different human influences will help us predict how organisms might respond to their rapidly changing environments.

## Acknowledgements

I wish to thank Marc-Olivier Beausoleil, Lara Bernasconi, Carlos Camacho, Sofia Carvajal, Angela Hansen, Célina Leuba, and Ashley Saulsberry for their assistance in the field. Permits were obtained with the assistance of Sofia Carvajal, Jaime Chaves, Jeff Podos, the Charles Darwin Research Centre, the Galápagos Science Centre, the Galápagos National Park, and the Universidad de San Francisco (Permits Nos PC-25-15, PC-42-16, PC-03-18). I thank Sarah Knutie for numerous helpful suggestions on ideas and methods, Ben Haller, Andrew Hendry, and Claire Spottiswoode for extensive comments and friendly reviews of the manuscript, and Tom Houslay for guidance with statistics. I thank Dan Blumstein and an anonymous reviewer for comments that greatly improved the manuscript. My research has been generously supported by the Natural Sciences and Engineering Research Council of Canada (Banting Postdoctoral Fellowship), Le Fonds Québécois de la Recherche sur la Nature et les Technologies (postdoctoral research fellowship), Clare Hall (Whitehead Fund), Christ’s College (Galápagos Islands Visiting Scholarship Scheme), Newnham College (Phyllis and Eileen Gibbs Travelling Research Fellowship), the British Ornithologists’ Union, and McGill University (Department of Biology Vineberg Fellowship). I have no conflicts to declare. Lastly, many thanks to Andrew Hendry, Jeff Podos, and Joost Raeymaekers for introducing me to the magical place that happened to be visited, some 185 years ago, by someone named Charles Darwin.

## Author’s contributions

KMG conceived and designed this research project, collected and analysed the data, and wrote the manuscript.

## Data Accessibility

Data will be archived on Dryad and on the author’s academic website.

