## Supplemental Files for "Human influences on antipredator behaviour in Darwin’s finches"

Supplemental Table 1. Summary and characteristics of the islands from which data were collected. Sample size indicates the number of finches chased on a specific island/population within island. Population estimates (Pop. N) are from 2015. Plus (**+**) and dash (**—**) indicates presence or absence of a given predator with cats and rats being invasive, and snakes, hawks, and owls being local and endemic predators.

| **Island** | **Status** | **Sample Size** | **Cats^2^** | **Rats^2^** | **Snakes^3^** | **Hawks^3^** | **Owls^3^** |
| --- | --- | --- | --- | --- | --- | --- | --- |
| Santa Fe | Pristine | 15 | **—** | **—** | **+** | **+** | **+** |
| Española | Pristine | 26 | -**—** | **—** | **+** | **+** | **+** |
| Floreana | Urban; Pop. N = 111^4^ | 18 | **+** | **+** | **+** | **—** | **+** |
|  | Non-Urban | 28 |  |  |  |  |  |
| Isabela | Urban; Pop. N = 2,164^4^ | 6 | **+** | **+** | **+** | **+** | **+** |
|  | Non-Urban | 12 |  |  |  |  |  |
| San Cristobal | Urban; Pop. N = 7,199^4^ | 12 | **+** | **+** | **+** | **—** | **+** |
|  | Non-Urban | 26 |  |  |  |  |  |
| Santa Cruz | Urban; Pop. N = 11,822^4^ | 27 | **+** | **+** | **+** | + | **+** |
|  | Non-Urban | 29 |  |  |  |  |  |
| Baltra | Eradicated 2003^5^ | 9 | **—** | **—** | **+** | **—** | **+** |
| North Seymour | Eradicated 2006^6^ | 19 | **—** | **—** | **+** | **—** | **+** |

^1^ Jackson, M. H. (2019). *Galapagos*. The University of Calgary Press, Calgary

^2^ Phillips, R.B., Wiedenfeld, D.A. & Snell, H.L. (2012). Current status of alien vertebrates in the Galápagos Islands: invasion history, distribution, and potential impacts. *Biol. Invasions*, 14, 461–480.

^3^ Swash, A. & Still, R. (2005). *Birds, Mammals, and Reptiles of the Galápagos Islands - An Identification Guide*. 2nd edn. Yale University Press, New Haven.

^4^ *Memoria Estadística Galápagos*. (2017). .

^5^ Phillips, R.B., Cooke, B.D., Campbell, K., Carrion, V., Marquez, C. & Snell, H.L. (2005). Eradicating feral cats to protect Galapagos Land Iguanas: methods and strategies. *Pacific Conserv. Biol.*, 11, 257–267.

^6^ Carrión, V., Sevilla, C. & Tapia, W. (2008). Management of introduced animals in Galapagos. *Galapagos Res.*, 65, 46–48.

Supplemental Figure 1. Map of the islands where FID was quantified for *Geospiza fuliginosa.* Four islands in orange have the presence of invasive predators and a permanent human population. Two islands in green are pristine with no presence of invasive predators, and two islands in purple are where invasive predators have been successfully eradicated.

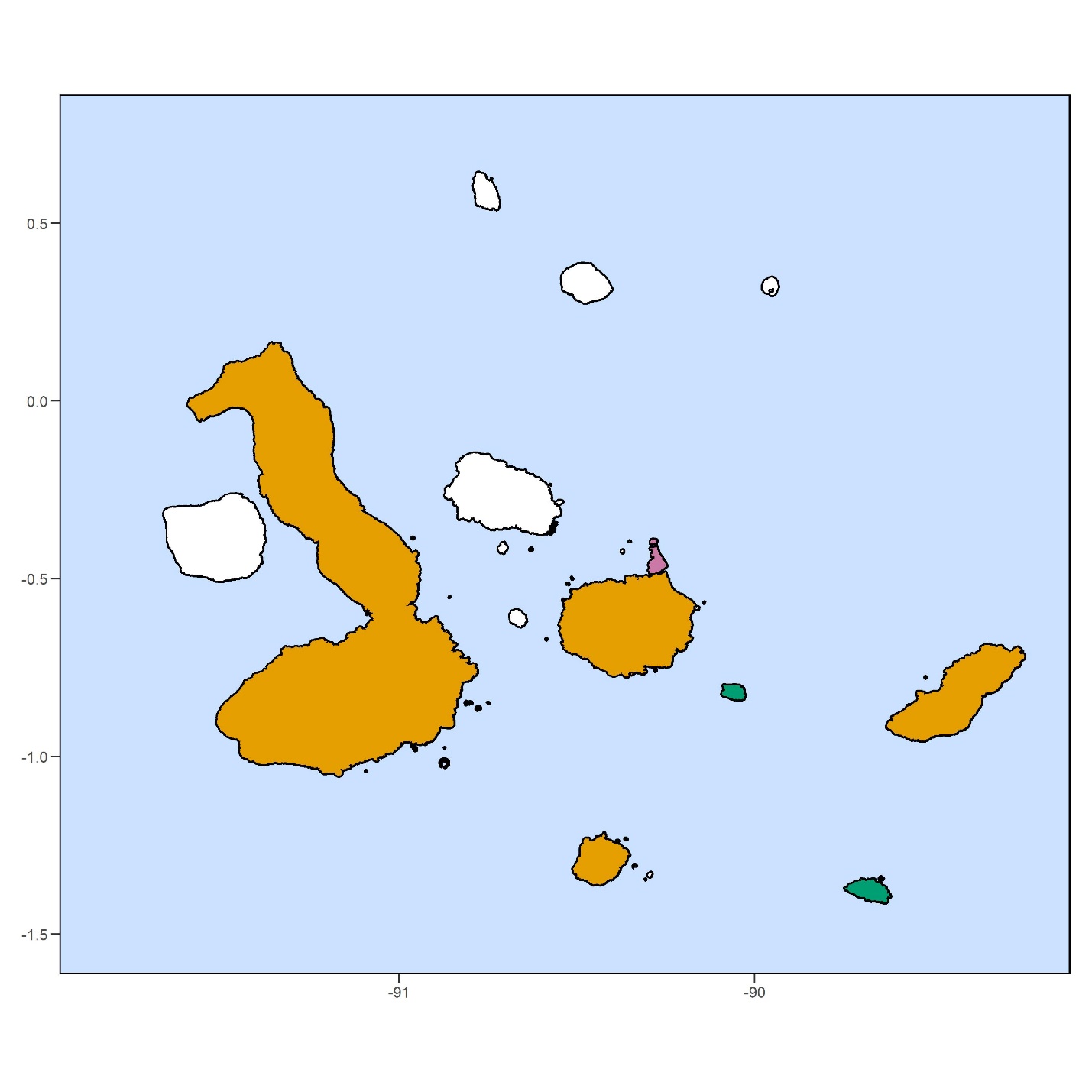

Supplemental Figure 2. FID in relation to starting distance for each island. Starting distance and island had a significant interaction in the full model looking at the effect of island on FID (Table 2). Data shown are from Baltra, North Seymour, Española, Santa Fe, and from non-urban sites on Floreana, Isabela, San Cristobal, and Santa Cruz. The interaction is driven by a negative relationship between FID and starting distance on San Cristobal. However, overall, starting distance was not correlated with FID (black regression line; Table 2)

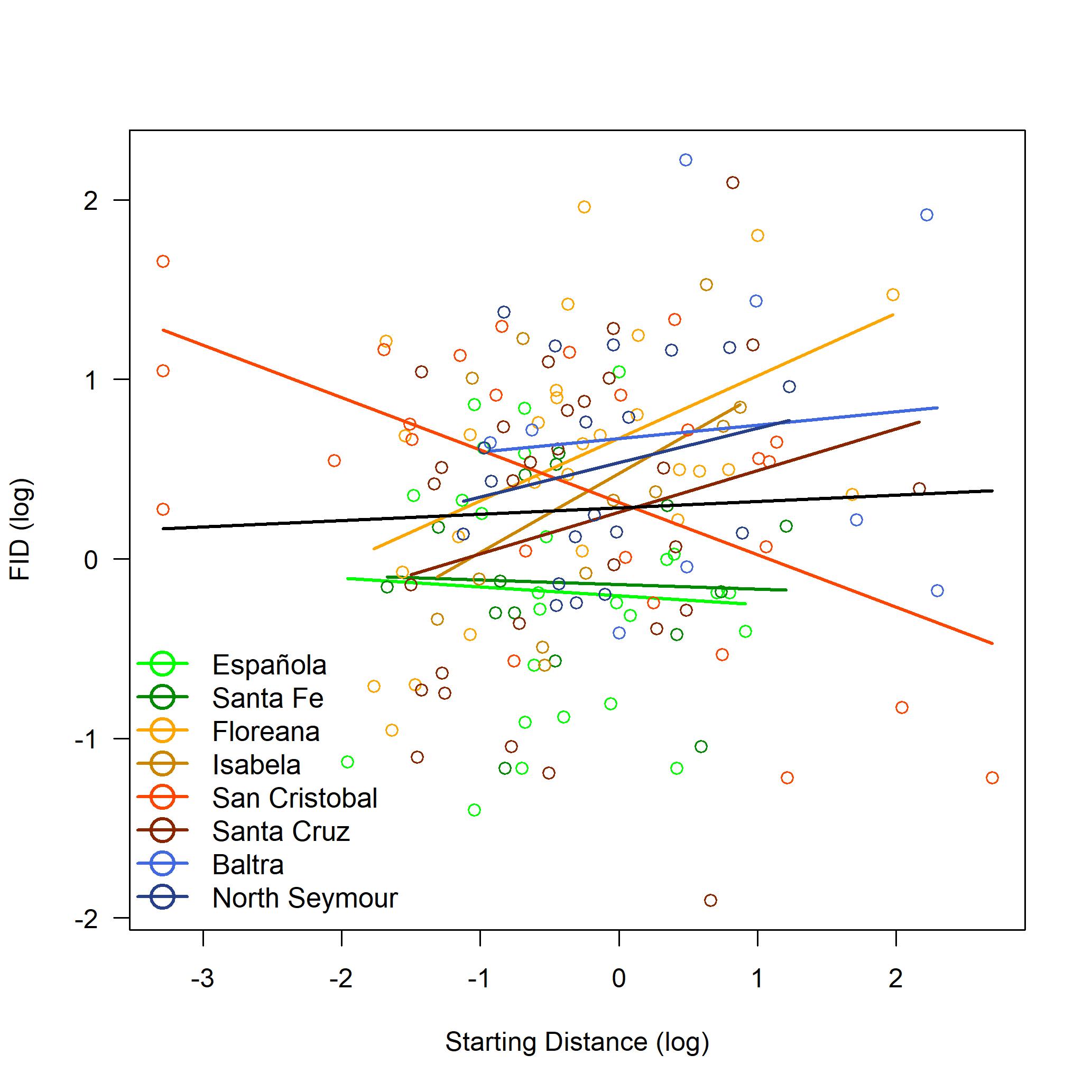

Supplemental Figure 3. FID in relation to group size. Data shown are from the four islands with human populations (Floreana, Isabela, San Cristobal, and Santa Cruz). In blue are data from urban sites and in pink are data from non-urban sites. FID increased with group size at both urban and non-urban sites.

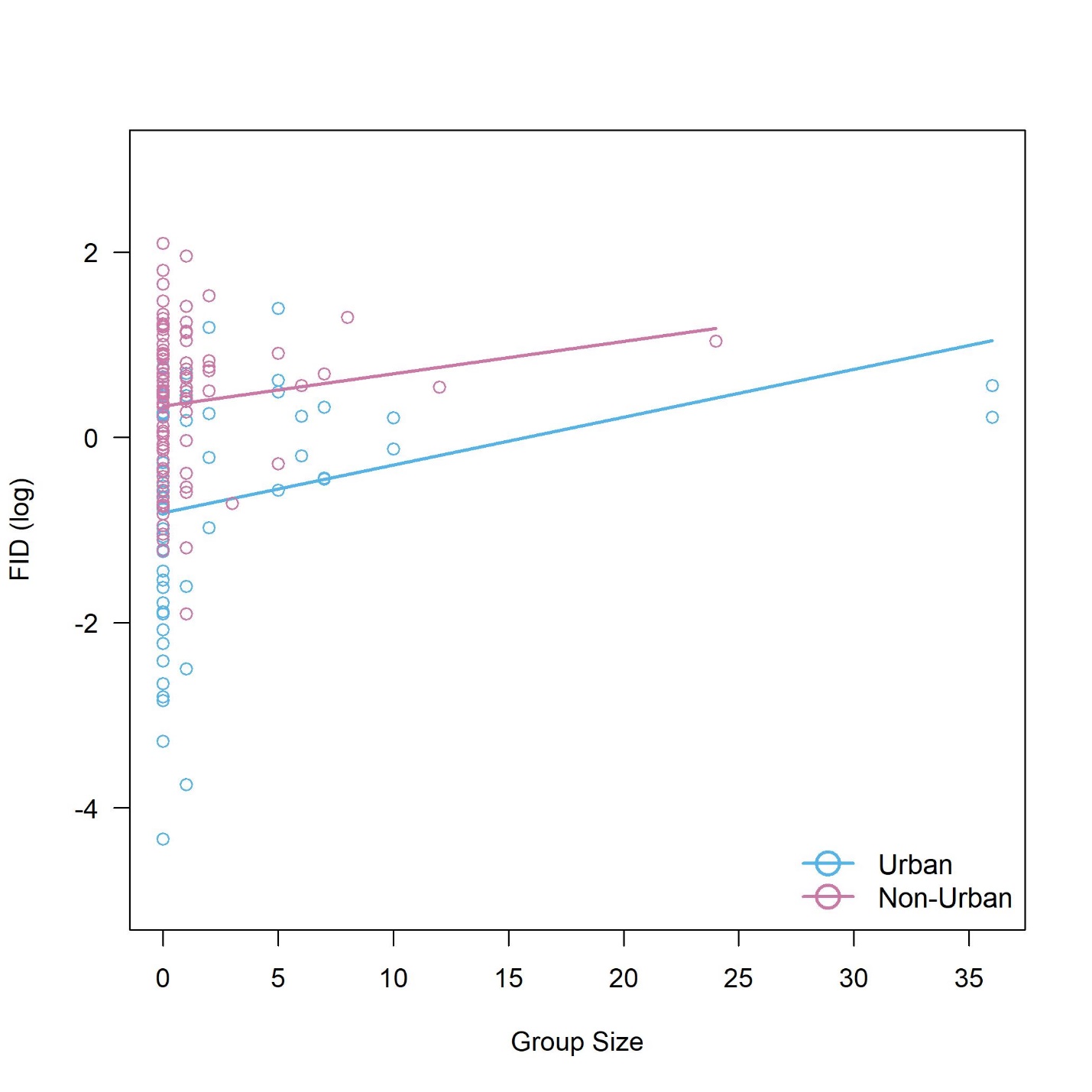
